# Exposure to low doses of *Batrachochytrium dendrobatidis* reveals variation in resistance in the Critically Endangered southern corroboree frog

**DOI:** 10.1101/2024.12.11.628040

**Authors:** Mikaeylah J. Davidson, Lee Berger, Amy Aquilina, Melissa Hernandez Poveda, Daniel Guinto, Michael McFadden, Deon Gilbert, Damian Goodall, Kyall R. Zenger, Lee F. Skerratt, Tiffany A. Kosch

**Affiliations:** Veterinary Biosciences, Faculty of Science, University of Melbourne, Werribee, VIC, Australia; Welfare, Conservation and Science, Taronga Conservation Society Australia, Mosman, NSW, Australia; Wildlife Conservation and Science, Zoos Victoria, Parkville, VIC, Australia; Centre for Sustainable Tropical Fisheries and Aquaculture, James Cook University, Townsville, Qld 4811, Australia; College of Science and Engineering, James Cook University, Townsville, Qld 4811, Australia

**Keywords:** Chytridiomycosis, *Pseudophryne corroboree*, Conservation, Captive breeding, Repeated experiments, Experimental variation, Amphibians, Threatened species

## Abstract

Chytridiomycosis poses a significant extinction threat to many amphibians, including the critically endangered southern corroboree frog (*Pseudophryne corroboree*). Captive breeding programs have become essential to maintain populations while effective long-term conservation strategies are developed. Understanding the variation in susceptibility to chytridiomycosis within this species is essential in exploring the potential for selective breeding to enhance disease resistance. In this study, we conducted a large-scale *Batrachochytrium dendrobatidis* (*Bd*) exposure experiment involving 972 juvenile *P. corroboree* selected to ensure a broad genetic representation of the species. Three replicate experiments were conducted under uniform conditions, to assess individual susceptibility and compare results across replicate experiments. Significant variation was observed within and between experiments, with individual survival rates ranging from 44-74% across experiments, influenced notably by the zoo in which frogs were bred. Remarkably, 21-47% of exposed frogs remained *Bd*-negative, suggesting potential innate resistance. Infection intensity correlated positively with body condition, in one experiment, while age and size showed inconsistent effects on survival and infection rates across experiments, but younger and smaller frogs were more susceptible to infection and had lower survival. Among frogs that became infected, none cleared infection, with most progressing to terminal stages within an average of 69 days (ranging from 33 to 97 days). However, a few individuals maintained stable infection loads without displaying clinical signs of chytridiomycosis. This observed phenotypic variation in *P. corroboree* responses to *Bd* highlights the potential for selective breeding to improve survival outcomes in this species. The dataset generated from this study will be instrumental in guiding breeding strategies that strengthen conservation efforts for this critically endangered species.

## 1. Introduction

One of the primary drivers of global amphibian declines is the spread of the amphibian chytrid fungus, *Batrachochytrium dendrobatidis* (*Bd*), which has caused devastating population collapses and extinctions (Scheele et al. 2019, Wren et al. 2024). Despite extensive research efforts over the past two decades, there remain few sustainable solutions to control or mitigate the impacts of chytridiomycosis in wild populations (Berger et al. 2024). Consequently, many species continue to decline and increasingly rely on captive breeding programs for survival (Harding et al. 2016). Without developing strategies to combat *Bd*, many of these species’ risk remaining in captivity indefinitely or facing eventual extinction in the wild.

Amphibian responses to *Bd* infection vary widely, reflecting their broad biological and ecological diversity. Different species and populations show a range of outcomes, shaped by differences in their ecology, environment, and genetic makeup (James et al. 2015, Lips 2016, Zamudio et al. 2020). Experimental exposures have been a key tool for understanding these responses, as they allow for controlled manipulation of specific factors. However, experimental methods often differ significantly across labs and studies, with factors like the *Bd* isolate, infectious dose, passage number, exposure method, and duration, as well as environmental variables such as temperature all impacting disease outcomes (Kumar et al. 2020). Such inconsistencies can introduce unintended variation, distorting responses and complicating cross-study comparisons. Additionally, small sample sizes and a lack of repeatability in many studies further limit our understanding of the mechanisms underlying variation in response to *Bd*.

While some species have shown signs of natural recovery from *Bd* (Newell et al. 2013, Knapp et al. 2016, Voyles et al. 2018), others, like the southern corroboree frog (*Pseudophryne corroboree*; traditionally known as “Gyack”), remain critically endangered. Unlike many tropical species that experienced rapid population crashes following the introduction of *Bd* to Australia (Berger et al. 1998, Scheele et al. 2017), *P. corroboree* exhibited a more gradual decline. This slower decline was likely influenced by more variable environmental conditions of its alpine habitat, which may be less conducive to *Bd* proliferation compared to the constantly humid and warmer rainforests. Additional factors such as their long lifespan (over 20 years in captivity; Davidson et al. 2025) may have slowed their rate of decline, and their solitary, terrestrial nesting behaviour (Pengilley 1973) may have reduced disease transmission.

Nevertheless, *P. corroboree* remains highly susceptible to *Bd,* exhibiting high mortality rates in captivity, and low survival after release to the wild (Hunter et al. 2010, Kosch et al. 2019, Davidson et al. 2025). With a lengthy maturation period of 4-6 years, low annual reproductive output of 16-38 eggs (McFadden et al. 2013), and high embryo mortality rates (Davidson et al. 2022), this species has low recruitment rates limiting their capacity for recovery. Therefore, despite ongoing conservation efforts, including captive breeding and reintroduction programs, *P. corroboree* remains dependent on human intervention to prevent extinction (Hunter et al. 2018). This is largely due to the existence of reservoir hosts that facilitate the persistence of the pathogen in the environment (Brannelly et al. 2018). Prior research has identified phenotypic and genetic variation in *P. corroboree*’s response to *Bd* (Kosch et al. 2019, Davidson et al. 2025), suggesting that selective breeding could potentially be used to increase their resistance to *Bd* and improve their long-term recovery efforts.

While selective breeding is well established in agriculture to enhance traits such as production efficiency and disease resistance, its use in conservation remains untested. A key reason for this is that selective breeding for specific traits contrasts with typical captive breeding programs for wildlife, which focus primarily on maintaining genetic diversity and preventing inbreeding depression (Allendorf et al. 2012, Kosch et al. 2022). Additionally, selective breeding requires substantial resources, including well-managed breeding populations, accurate phenotypic data, and extensive genetic information, which can be challenging to obtain in conservation contexts. *P. corroboree* offers a unique opportunity to explore the efficacy of selective breeding in conservation, as this iconic species is intensively managed in captivity and comprehensive genomic resources have been developed, including a chromosome-level genome (#GCA_028390025.1) and a high-density SNP array (Davidson et al. 2024). However, to develop genomic predictions that inform selective breeding strategies, it is essential to first develop a comprehensive understanding of their phenotypic responses to *Bd*.

To investigate phenotypic variation in response to chytridiomycosis in the Critically Endangered, captively managed *P. corroboree*, we conducted a large-scale experimental *Bd* exposure experiment on frogs bred from founder animals. This experiment involved 972 juvenile frogs, encompassing a broad genetic representation of the species diversity in captivity. The experiment was replicated three times to enhance the robustness of our findings across time and experiments. Under relatively uniform conditions, we aimed to assess how *P. corroboree* responds to *Bd* exposure, thus providing valuable insights into susceptibility and resistance mechanisms. We hypothesised that there would be varying levels in the response to *Bd* in *P. corroboree*, with some frogs and families exhibiting greater resilience compared to others, and that the effects of experimental variables would remain consistent across experimental replications. The phenotypic dataset generated in this study is one of the most comprehensive to date for studying the effects of *Bd* on a threatened amphibian species. It will be instrumental in informing future work aimed at enhancing chytrid resistance, a key factors in ensuring the long-term conservation of this species. This study not only enhances our knowledge of *Bd* susceptibility in this species but represents a step forward in integrating genetic approaches into amphibian conservation strategies.

## 2. Methods

### 2.1. Animal husbandry

Over two years, a total of 972 captive-bred *P. corroboree* were obtained from Melbourne Zoo (MZ) (2021 N = 146 and 2022 N = 360) and Taronga Conservation Society Australia (TZ) (2021 N = 163 and 2022 N = 303). As captive breeding of *P. corroboree* occurs in group enclosures, full parentage assignment remains undetermined for most clutches. It was predicated that 54 full-sibling clutches were laid, ranging from 2 to 64 individuals each, with potential contributions from 59 sires and 99 dams representing the 15 historical *P. corroboree* populations. Efforts were made to establish approximately 50 clutches with around 20 offspring each to capture within- and between-family variation in traits under controlled experimental conditions. This approach ensured genetic representation from each historical population, allowed for the replication of sires across the two breeding years, and was within the zoos’ capacity to feasibly breed and raise offspring.

Due to logistical challenges, the experimental exposures were divided into three consecutive experiments, designated as experiments (Exp) 1-3. Frogs bred in 2021 were used exclusively in Exp 1, while those bred in 2022 were split equally by clutch between Exp 2 and 3. All protocols and procedures were consistent across experiments unless otherwise stated. Prior to experiments, frogs were housed in their respective clutches in group enclosures with gravel and live plants, misted daily with carbon-filtered water, and fed *ad libitum* three times weekly with springtails (*Collembola* sp.) and vitamin-dusted crickets (*Acheta domestica*). Two weeks before the experiments, frogs were transferred into individual housing with damp paper towels, with the experimental husbandry methods following those outlined in Davidson et al. (2025). Each frog was randomly assigned a tank number using a random number generator, with one animal from each clutch assigned to the control group. Across the experiments, the main factor that varied was room temperature. In Exp 1, the average temperature was 17.0°C (range: 14.8–20.6°C), compared with 16.8°C (range: 14.7–19.6°C) in Exp 2, and 16.6°C (range: 14.3–20.1°C) in Exp 3. Additionally, variations in animal availability led to differences in the timing of the experiments, resulting in discrepancies in both the age (measured as days since metamorphosis) and the size of the frogs at the start of each experiment. Overall, frogs were between 4.5-12 months old (mean = 9 ± 1.7 months), weighed between 0.28-1.52 g (mean = 0.94 ± 0.23 g), and their snout-vent lengths were from 15.4-28.6 mm (mean = 23.1 ± 2.1 mm) (Figure S1 and Table S1 for individual experiment metrics).

### 2.2. Experimental Bd exposures

We inoculated frogs with the *Bd* isolate Wastepoint-L. verreauxii-2013-LB-RW, which was collected from a site near Jindabyne, New South Wales, Australia. This site is near the former distribution of *P. corroboree* (see Davidson et al. (2025) Figure S1 for map). For each experiment, cryopreserved cultures were defrosted following Boyle et al. (2003) at passage six, and used at passage seven or eight.

*Bd* zoospores were quantified using a haemocytometer after being harvested from four-day-old cultures grown on TGhL agar plates. The plates were rinsed and then flooded with 2 mL of sterile Milli-Q water for 20 minutes before the suspension was passed through a 10 µm filter (Pluriselect). Frogs were inoculated with 2,000 zoospores in 3 ml water via a six-hour bath exposure in 70 ml containers, exposed by dripping inoculum into the container. The dose was chosen to maximise individual variability and was determined through a pilot study (Davidson et al. 2025). Control frogs (overall N = 54, Table 1) were inoculated using the same method but with inoculum from *Bd*-negative plates.

**Table 1.** Overview of the number of *P. corroboree* used in each experiment.

|  | No.<br>frogs | Frogs per zoo | No.<br>clutches | No.<br>controls | No.<br>exposed to<br>*Bd* |
| --- | --- | --- | --- | --- | --- |
| Exp 1 | 309 | MZ = 146<br>TZ = 163 | 20 | 20 | 289 |
| Exp 2 | 333 | MZ = 180<br>TZ = 153 | 34 | 17 | 316 |
| Exp 3 | 330 | MZ = 180<br>TZ = 150 |  | 17 | 313 |
| Overall | 972 | MZ = 506<br>TZ = 466 | 54 | 54 | 918 |

Note: The number of frogs for each experiment (exp) sourced from each zoo–Melbourne Zoo (MZ) or Taronga Zoo (TZ)–along with the breakdown of clutches, control frogs, and those exposed to *Bd*.

Following exposure, frogs were monitored daily for clinical signs of chytridiomycosis, and those exhibiting terminal signs–typified by a delay in the ability to right themselves–were euthanised in a buffered 0.2% MS-222 bath. Frogs were swabbed weekly to test for *Bd* as adapted and described in Davidson et al. (2025). Briefly, a standard five-stroke swabbing method was used on their venter, along each ventral side, and on each thigh and foot (Brannelly et al. 2015a). DNA was extracted using Prepman Ultra (Applied Biosystems), with bead beating (Brannelly et al. 2015b). Quantitative PCR (qPCR) was conducted in singlicate for each sample, using seven *Bd* plasmid standards (Pisces Molecular) (Boyle et al. 2004, Brannelly et al. 2020).

### 2.3. Statistical analysis

All statistical analyses were conducted in R (4.1.2) using the RStudio interface (RStudio Team 2020, R Core Team 2021). Since all 54 controls remained healthy and uninfected, they were excluded from the analysis.

#### 2.3.1. Individual experiments

We compared frog survival between clutches by Cox regression using the *survival* package (Therneau & Lumley 2015). Separate regression models were constructed for zoo, age of the frog, and body condition (log mass / log snout-venter length), which were included as covariates to assess the effect on survival rates. All models were clustered by clutch to account for potential within-clutch correlations in survival rates. This approach allowed for a robust analysis of how different factors may influence survival outcomes in *P. corroboree*, considering both individual and environmental variations.

Infection loads were converted from Internal Transcribed Spacer (ITS1) copy number (Davidson et al. 2025) to zoospore equivalents (ZE) and log10 transformed before linear mixed model analysis with lme4 (Bates et al. 2015). We used *emmeans* (Lenth 2022) to obtain and compare the rate of increase in infection load from the model. All models included the fixed effect of week as a continuous variable and the random effect of frog ID. A random slope of week for each frog was also included.

To evaluate the factors influencing infection load dynamics in *P. corroboree*, we employed a series of four statistical models. The first model compared the rate of increase in infection load between clutches, by treating clutch as a fixed effect and including an interaction term of clutch and week. The second model compared the effect of zoo on infection dynamics using zoo as a fixed effect, an interaction term between zoo and week, and a random effect of clutch with a random slope of week for each clutch. The third model examined the effect of age on the rate of infection load increase, with frog age at the start of the experiment as a fixed effect and an interaction term between age and week, including a random effect of clutch. The fourth model focused on the effect of body condition on the rate of infection load increase, using body condition as a fixed effect, an interaction term between body condition and week, and a random effect of clutch. Frogs that remained *Bd* negative for the entire experiment were excluded from the models (Table 2).

**Table 2.**
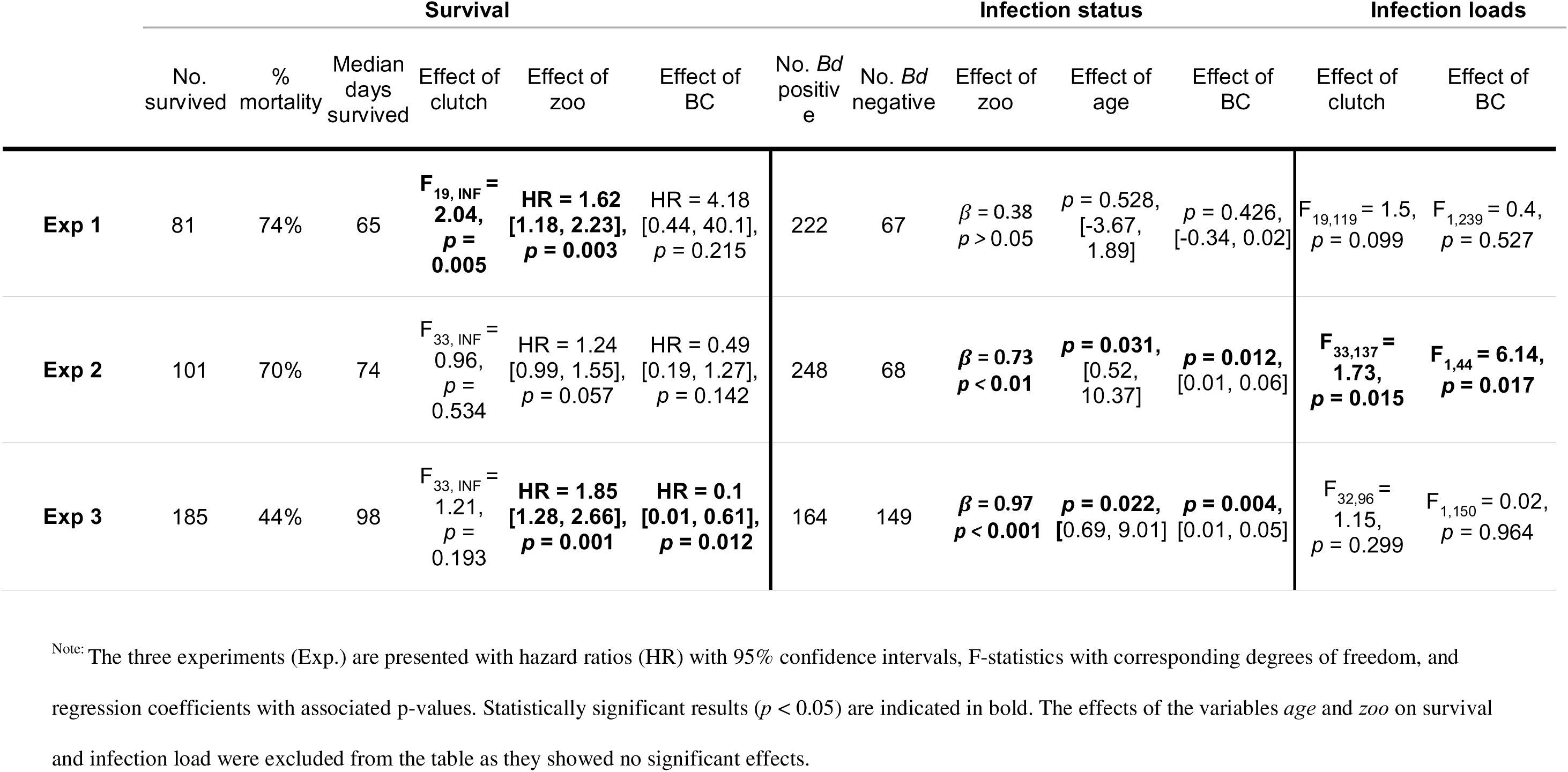
Summary of survival, infection status, and infection loads for *P. corroboree* across individual experiments.

#### 2.3.2. Combined analysis

A random-effects meta-analysis (REM) was used to pool the results from the three individual experiments using the restricted maximum likelihood method (Balduzzi et al. 2019). This method accounts for potential heterogeneity across the three experiments by considering multiple factors that could introduce variability. These factors include the year the frogs were bred, the zoo where they were bred, their age and size, imprecision in infection methods, the time of year the experiment was conducted and other unmeasured environmental effects.

### 2.4. Ethics & data availability

All procedures were conducted under approval of the University of Melbourne’s Animal Ethics Committee (Application 2021-22144-24454-5), and Wildlife Act 1975 Research Authorisation permit number 10010261. All data and code required to conduct the analysis are available at https://figshare.com/s/89fe2c542c884870479a.

## 3. Results

### 3.1. Individual experiments

The overall mortality rates among the *Bd-*exposed frogs across the three experiments were 74%, 70% and 44% (Exp 1-3 respectively) (Table 2 and Figure 1A). The number of frogs that survived to the end of the experiment, the median survival time, and the number of *Bd-* negative frogs also differed across the three experiments (Table 2). All 54 controls across the three experiments remained uninfected and survived the 98-day experiments.

**Figure 1.**
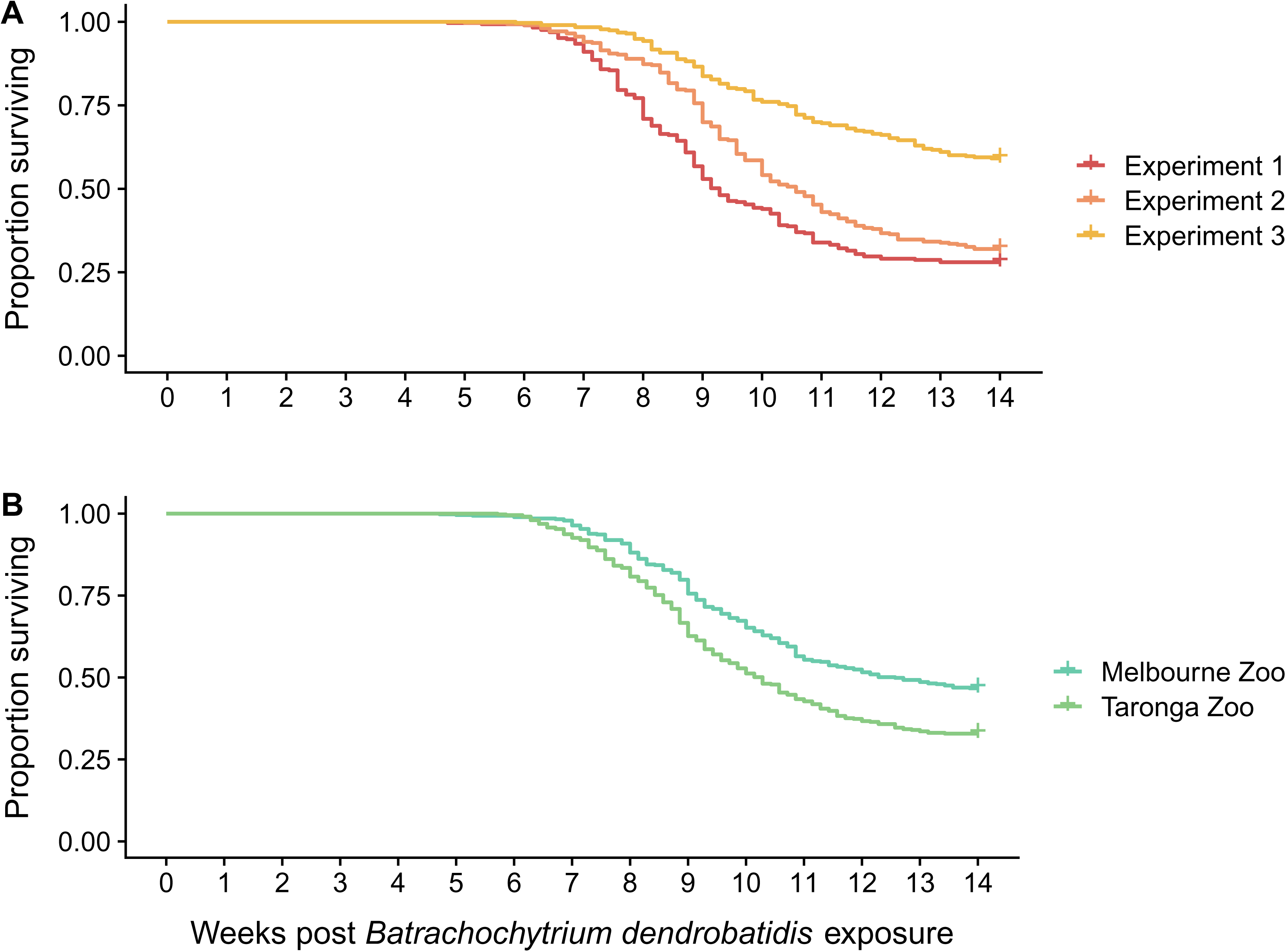
Survival outcomes of *P. corroboree* exposed to *Bd* across (A) the three individual experiments and (B) the respective zoo where they were bred and raised.

The zoo which frogs originated from significantly impacted their survival times in Exp 1 and 3, with frogs from TZ exhibiting an increased risk of mortality; however, this effect was not observed in Exp 2 (Table 2 and Figure S2). Additionally, there was a difference in survival time between clutches in Exp 1, but no difference was detected in Exp 2 or 3 (Table 2). In Exp 3, the body condition of the frogs significantly impacted survival time, with smaller frogs having a higher risk of mortality. This effect, however, was not observed in Exp 1 or 2 (Table 2). Lastly, age did not influence survival across any of the experiments.

The infection status of frogs after exposure (*Bd*-positive vs. *Bd*-negative) was significantly influenced by the frog’s origin, age, and body condition in Exp 2 and 3, but not in Exp 1 (Table 2, Figure S3 and S4). Frogs from TZ, as well as younger and smaller individuals, were more likely to become infected compared to frogs from MZ, which were older or larger. This difference may be attributed to significant differences in age, length, and weight between frogs from MZ and TZ in Exp 2 and 3 (Table S2 and Figure S5).

The rate of increase in infection loads varied significantly between clutches and was influenced by body condition in Exp 2; however, no differences were observed in Exp 1 or 3. Additionally, neither zoo nor age had an effect on infection loads in any of the experiments (Figure 2).

**Figure 2.**
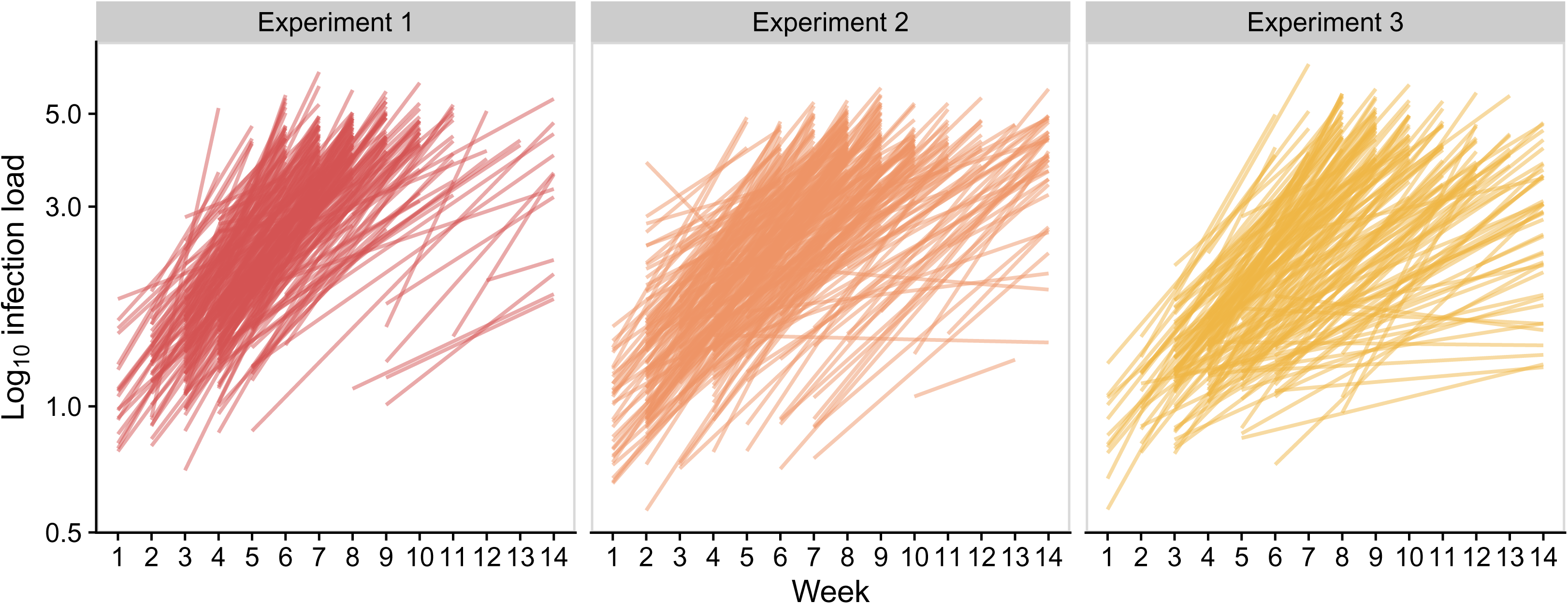
*Bd* infection loads for each individual across the three experiments. Frogs that remained *Bd-*negative have been omitted (N = 67, 68, 149, respectively).

### 3.2. Combined analysis

Across the three experiments, *Bd* exposure resulted in an overall mortality rate of 60% (551 out of 918 frogs). Consequently, 367 Bd-exposed frogs survived the 14-week study period, with a median survival time of 77 days. Of these 367 surviving frogs, 284 remained *Bd*-negative for the entire duration of the experiments.

Survival time was significantly influenced by the zoo the frogs originated from, with frogs from TZ exhibiting increased mortality rates across experiments (HR: 1.52 95% CI [1.19, 1.82], *p* < 0.001, I^2^ = 44%) (Table S3 and Figure 1B). Additionally, the age and size of the frogs did not significantly impact survival times in the REM, with the direction of effect differing across experiments. In Exp 1 younger and smaller frogs had higher survival rates, whereas in Exp 2 and 3 younger and smaller frogs experienced increased mortality (Table S3).

The rate of infection load increase showed a significant, albeit weak, positive correlation with the body condition of frogs (0.176, 95% CI [0.006, 0.347], *p* = 0.043, I² = 0%). Larger frogs exhibited increased infection loads in Exp 1 and 2, while Exp 3 showed a negligible association between body condition and infection load (Table S4). Although age did not have a significant overall effect on infection rates, a similar trend was observed, with older frogs in Exp 1 and 2 having steeper increases in infection loads, compared to younger frogs in Exp 3. Additionally, the zoo of origin did not significantly impact infection loads, but frogs from TZ exhibited steeper increases in infection loads in Exp 1 and 3 compared to Exp 2 where MZ frogs had higher rates of infection (Table S4).

## 4. Discussion

By performing the largest experimental *Bd* challenge study to date, using 972 juvenile *P. corroboree* bred to represent the genetic breadth of the species, we detected phenotypic variation in disease resistance that might have gone unnoticed with smaller sample sizes. Evidence from repeated experiments showed *P. corroboree* susceptibility varied between individuals and there were differences in phenotypic responses between exposures, with survival rates ranging from 26-56 % across the three experiments. Interestingly, 21-47 % of *Bd*-exposed frogs remained uninfected, explaining the moderate overall mortality rates.

Among those that became infected, most individuals reached terminal stages within 69 days on average (ranging from 33 to 97 days), although a few individuals maintained stable infection loads without exhibiting clinical signs of chytridiomycosis. Additionally, no single experimental variable had a consistent effect on survival or infection across all experiments, suggestive of different factors, potentially including some not measured here such as genetics, influencing outcomes in each experiment. Thus, our large and repeated experiments provided a unique opportunity to examine variation in response to *Bd* by using the same species under uniform conditions.

As part of a broader initiative to enhance chytrid resistance in *P. corroboree* through selective breeding (Kosch et al. 2019, Davidson et al. 2024, Davidson et al. 2025), this study aimed to collect and analyse phenotypic responses to *Bd* infection. Although ideally this study would have been conducted as a single experiment to minimise variation in factors beyond individual resistance, logistical challenges necessitated conducting the research as three separate experiments. Despite our efforts to minimise variation–by conducting the experiments under identical conditions, having the same person collect all data, and adhering to consistent protocols for husbandry and molecular work–inherent variation resulting from splitting the experiments was beyond our control. As a counter to that, replication of experiments across time increased the likelihood of distinguishing the key factors influencing the outcome of *Bd* exposure.

We observed variation in both survival and infection rates across the experiments, highlighting the diverse responses of *P. corroboree* to *Bd*. Notably, nearly one-third of the exposed frogs remained negative for *Bd*, which contrasts with previous laboratory exposures for this species resulting in infection rates of 98–100 % (Kosch et al. 2019, Davidson et al. 2025). This past work involved exposing adults to 1 million zoospores, and juveniles to a range of high to low doses. However, our lower infection rates align with infection dynamics observed in the wild, where prevalence ranged from 44-60 % (Hunter et al. 2010). The prevalence observed in the wild may have contributed to the slow decline of *P. corroboree* and may be due to lower exposure doses. Because *P. corroboree* are fully terrestrial and do not congregate for breeding, their exposure to *Bd* in the wild may occur at lower frequencies. Additionally, the alpine habitat historically occupied by *P. corroboree* is diverse, encompassing a range of environmental niches and fluctuations in hydrology and temperature, which could further contribute to lower rates of exposure and infection. Previous work in juvenile *P. corroboree* demonstrated that exposure to low infectious doses allows for variation in responses to *Bd* to be detected (Davidson et al. 2025). Therefore, the low exposure dose used in our study may more accurately reflect natural exposures and explain the variation in infection levels observed in the wild. However, the constant temperature and humidity in our experimental tanks were likely more favourable for disease progression than the natural habitat of *P. corroboree*.

The observed variability in *Bd*-positive frogs in our study could be attributed to the low infectious dose combined with the indirect method of experimental *Bd* exposure used. As noted earlier, 31 % of frogs never tested positive for *Bd*, potentially due to some individuals not having been adequately exposed, while others might have resisted low exposure through an innate cutaneous response (Woodhams et al. 2007, Burkart et al. 2017, Grogan et al. 2018). Although a moderate proportion of frogs remained *Bd*-negative across all three experiments, none that tested positive cleared their infection. The relatively long incubation periods suggest that the low dose and experimental conditions enabled a host response in some individuals that contributed to their slow disease progression. But ultimately the lack of clearance suggests *P. corroboree* were unable to mount an effective adaptive immune response, even though some individuals reached a plateau in their infection load. This contrasts with the findings of Kosch et al. (2019), who documented clearance in a small subset of adult *P. corroboree*.

The zoo where the frogs were bred was a significant source of variation in our study. *P. corroboree* from TZ exhibited a higher mortality risk compared to those from MZ. This disparity may stem from differences in the genetic representation of founding frogs across the two zoos, as well as variations in the breeding conditions or environmental factors associated with each zoo. Future work should compare the genome-wide genetic diversity of the founders and their offspring, including the animals used here, across the two zoos. Such genetic assessments are particularly relevant given that previous studies have shown increased heterozygosity in the major histocompatibility complex (MHC) to be associated with improved survival in several species (Savage & Zamudio 2011, Bataille et al. 2015), including *P. corroboree* (Kosch et al. 2019). Since many traits linked to disease resistance are polygenic, genome-wide heterozygosity may be an important factor in predicting and increasing the resilience of *P. corroboree* to *Bd*.

The health and resilience of the captive-bred frogs produced by the two zoos may also be influenced by differences in their husbandry practices. Variations in protocols for key husbandry aspects such as parental care, overwintering, tadpole rearing, and diet all likely contributed to the observed differences in age and size among the frogs. While age and size did not consistently impact survival, younger and smaller frogs in Exp 2 and 3–originating from TZ–had an increased likelihood of infection. Conversely, in Exp 1, older and larger frogs from the same zoo showed reduced survival. This suggests that the zoo where frogs originated from may be a stronger predictor of *P. corroboree* susceptibility than individual factors like age or size.

Interestingly, while smaller frogs were more likely to become infected, it was the larger frogs that exhibited higher infection intensities once infected. This positive correlation between body condition and infection load contrasts with findings from other studies, where smaller frogs typically have increased infection loads (Gervasi et al. 2017, Wu et al. 2018, Meurling et al. 2024). Despite higher infection intensities in larger frogs, body condition only influenced survival in one of the three experiments, where smaller frogs had lower survival. Consequently, body condition and infection loads in *P. corroboree* did not consistently influence survival across the experiments. This suggests that while body size may play a role in infection dynamics, its relationship with survival is complex and may depend on other factors, including immune function and rearing environment.

Experimental *Bd* exposures are commonly used to assess species susceptibility; however, large or repeated experiments are rare, likely due to the practical and ethical challenges associated with working with threatened species. Our findings demonstrate that, while experimental outcomes can vary even under controlled conditions, the combined results are generally consistent, in line with previous studies (Carvalho et al. 2023). Nevertheless, our repeated experiments highlight that while broader trends can be identified, individual experiments may over or underrepresent the effects of certain variables. Most of the variation we observed was linked to the zoo in which frogs were bred and raised. Therefore, relying on a single experiment, particularly from a single captive colony or potentially wild population could skew disease outcomes and fail to capture the full range of response to *Bd*. While variation between experiments should be minimised, within experiment variability, particularly related to the animals themselves, can lead to more robust and representative findings. Since our aim was to study phenotypic variation in response to *Bd*, variability alongside large sample sizes provided differential outcomes in response to exposure.

In conclusion, we show that southern corroboree frogs display significant phenotypic variation in response to *Bd* under controlled experimental conditions. Although many frogs exhibited high susceptibility to infection, evidence suggests that a portion of the population possesses the ability to either resist or tolerate infection without developing clinical chytridiomycosis, provided that the level of *Bd* exposure is low enough. Since all frogs used in this study were captive-bred and the experiments were conducted under uniform conditions, an important next step will be to compare results with frogs raised in outdoor mesocosms. By comparing laboratory exposures with mesocosm conditions–where frogs are subject to changing weather conditions, have microhabitat choice, and a natural diet–we can gain insights into the ecological factors that influence *P. corroboree* susceptibility to *Bd*.

Given that a large proportion of frogs were resistant to infection, future research should also investigate the early innate immune response in *P. corroboree*, and whether susceptibility and immune responses vary across different age classes. Furthermore, we plan to leverage this dataset to explore the genetic associations of disease resistance within this population. Such insights will be instrumental in developing selective breeding programs aimed at increasing the resilience of *P. corroboree* against *Bd*. The slow declines observed in *P. corroboree*, coupled with the resistance shown by some individuals, suggest that even moderate improvements in disease resistance could facilitate recovery efforts for this critically endangered species, ultimately increasing their chances of survival in the wild.

## 5. Acknowledgements

We express our gratitude to David Hunter and the *Pseudophryne corroboree* keepers at Melbourne Zoo and Taronga Conservation Society, as well as everyone who has been involved in their success across the other partner zoos and programs. Special thanks to Carley Gringer, Jasper White, Katie Lee, Mandy To, and Quinn Higgs for their assistance with sample processing and animal husbandry, and Cameron Patrick from the Statistical Consulting Centre at the University of Melbourne for statistical guidance. This study was funded by Australian Research Council grants (FT190100462 to LFS and LP200301370 to LFS, LB, KRZ, DG and MM), and a University of Melbourne Graduate Research Scholarship was awarded to MJD.

