## Supplementary material for "Exposure to low doses of *Batrachochytrium dendrobatidis* reveals variation in resistance in the Critically Endangered southern corroboree frog": Table S; Figure S

**Table S1**. Summary of demographic and physical characteristics of the *P. corroboree* used across the three *Bd* infection experiments.

| Year bred | Experiment | MZ | TZ | Total No. | No. clutches | Age (days) | Weight (g) | SVL (mm) |
|---|---|---|---|---|---|---|---|---|
| 2021 | 1 | 146 | 163 | 309 | 20 | 269 - 303 | 0.28 - 1.40 | 18.6 – 26.0 |
| 2022 | 2 | 180 | 153 | 333 | 34 | 146 - 242 | 0.35 - 1.44 | 16.9 - 27.3 |
| | 3 | 180 | 150 | 330 | | 261 - 357 | 0.27 - 1.52 | 15.4 - 28.6 |

Abbreviations: MZ, Melbourne Zoo; TZ, Taronga Zoo; SVL, snout-venter length.

**Table S2.** Comparison of frog age and morphometrics at the start of the three experiments, between the frogs bred across the two zoos.

| Effect | Experiment | t-value | Degrees of Freedom | p-value | MZ | TZ | 95% Confidence Interval |
|---|---|---|---|---|---|---|---|
| | | | | | Mean age (days) | | |
| Age | 1 | -21.367 | 242.44 | < 0.001 | 275 | 292 | -18.20, -15.13 |
| | 2 | 22.55 | 261.82 | < 0.001 | 202 | 172 | 27.60, 32.89 |
| | 3 | 22.34 | 251.94 | < 0.001 | 317 | 287 | 27.02, 32.25 |
| | | | | | Mean SVL (mm) | | |
| SVL | 1 | -3.86 | 286.8 | < 0.001 | 22.9 | 23.4 | -0.74, -0.24 |
| | 2 | 30.29 | 298.43 | < 0.001 | 24.3 | 20.2 | 3.79, 4.31 |
| | 3 | 15.89 | 304.5 | < 0.001 | 25.3 | 22.1 | 2.73, 3.49 |
| | | | | | Mean weight (g) | | |
| Weight | 1 | 0.085 | 280.83 | 0.933 | 0.995 | 0.993 | -0.032, 0.04 |
| | 2 | 37.67 | 312.52 | < 0.001 | 1.102 | 0.610 | 0.47, 0.52 |
| | 3 | 15.59 | 284.98 | < 0.001 | 1.096 | 0.789 | 0.27, 0.35 |

Abbreviations: MZ, Melbourne Zoo; TZ, Taronga Zoo; SVL, snout-venter length.

**Table S3.** Summary of the effects of zoo, age, and body condition on the survival of *P. corroboree* across the three *Bd* infection experiments.

| Effect | Experiment | Effect Estimate | Hazard Ratio | p-value | $I^2$ |
|---|---|---|---|---|---|
| Zoo | **1** | 0.483 | 1.620 [1.229, 2.141] | | |
| | **2** | 0.215 | 1.239 [1.056, 1.261] | | |
| | **3** | 0.613 | 1.845 [1.298, 2.606] | | |
| | **REM** | 0.419 | 1.520 [1.198, 1.819] | < **0.001** | 44.20% |
| Age | **1** | 0.009 | 1.009 [0.9968, 1.0217] | | |
| | **2** | -0.005 | 0.995 [0.9885, 1.0025] | | |
| | **3** | -0.010 | 0.990 [0.9809, 1.0000] | | |
| | **REM** | -0.002 | 0.998 [0.9879, 1.0074] | 0.625 | 64.60% |
| Body condition | **1** | 1.429 | 4.169 [0.382, 46.35] | | |
| | **2** | -0.714 | 0.490 [0.0015, 0.624] | | |
| | **3** | -2.296 | 0.101 [0.0006, 0.677] | | |
| | **REM** | -0.645 | 0.525 [0.0040, 3.414] | 0.500 | 64.90% |

Note: The random effects model (REM) provides a combined estimate across all experiments. Hazard ratios are shown with their 95% confidence intervals. $I^2$ values represent the percentage of variation due to heterogeneity among experiments.

**Table S4.** Summary of the effects of zoo, age, and body condition on infection loads of *P. corroboree* across the three *Bd* infection experiments.

| Effect | Experiment | Effect Estimate | p-value | $I^2$ |
|---|---|---|---|---|
| Zoo | 1 | 0.036 [-0.0267, 0.0987] | | |
| | 2 | -0.040 [-0.0811, 0.0013] | | |
| | 3 | 0.020 [-0.0412, 0.0804] | | |
| | REM | 0.0003 [-0.0485, 0.0491] | 0.989 | 59.90% |
| Age | 1 | 0.0002 [-0.0029, 0.0032] | | |
| | 2 | 0.001 [-0.0004, 0.0020] | | |
| | 3 | -0.001 [-0.0023, 0.0009] | | |
| | REM | 0.0002 [-0.0011, 0.0014] | 0.794 | 12.00% |
| Body condition | 1 | 0.191 [-0.3933, 0.7752] | | |
| | 2 | 0.237 [0.0548, 0.4186] | | |
| | 3 | -0.008 [-0.3484, 0.3322] | | |
| | REM | 0.176 [0.0056, 0.3465] | **0.043** | 0.00% |

Note: The random effects model (REM) provides a combined estimate across all experiments. The regression coefficient (effect estimate) is shown with their 95% confidence intervals. $I^2$ values represent the percentage of variation due to heterogeneity among experiments.

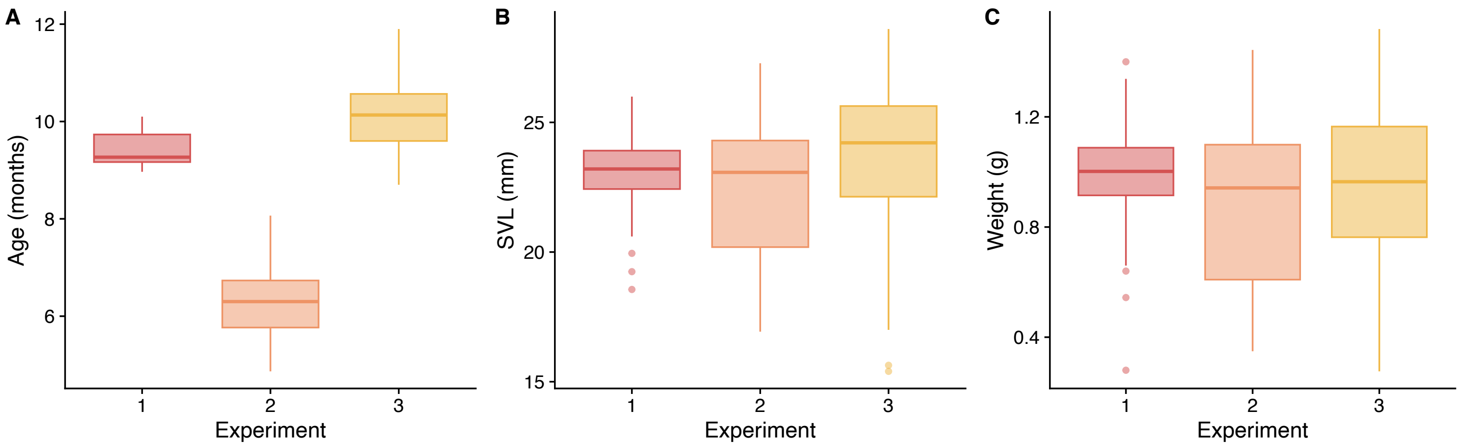


**Figure S1.** Comparison of (**A**) age; (**B**) snout-vent length (SVL); and (**C**) weight of the 972 *P. corroboree* at exposure for the three experiments (N = 309, 333, 330, respectively).

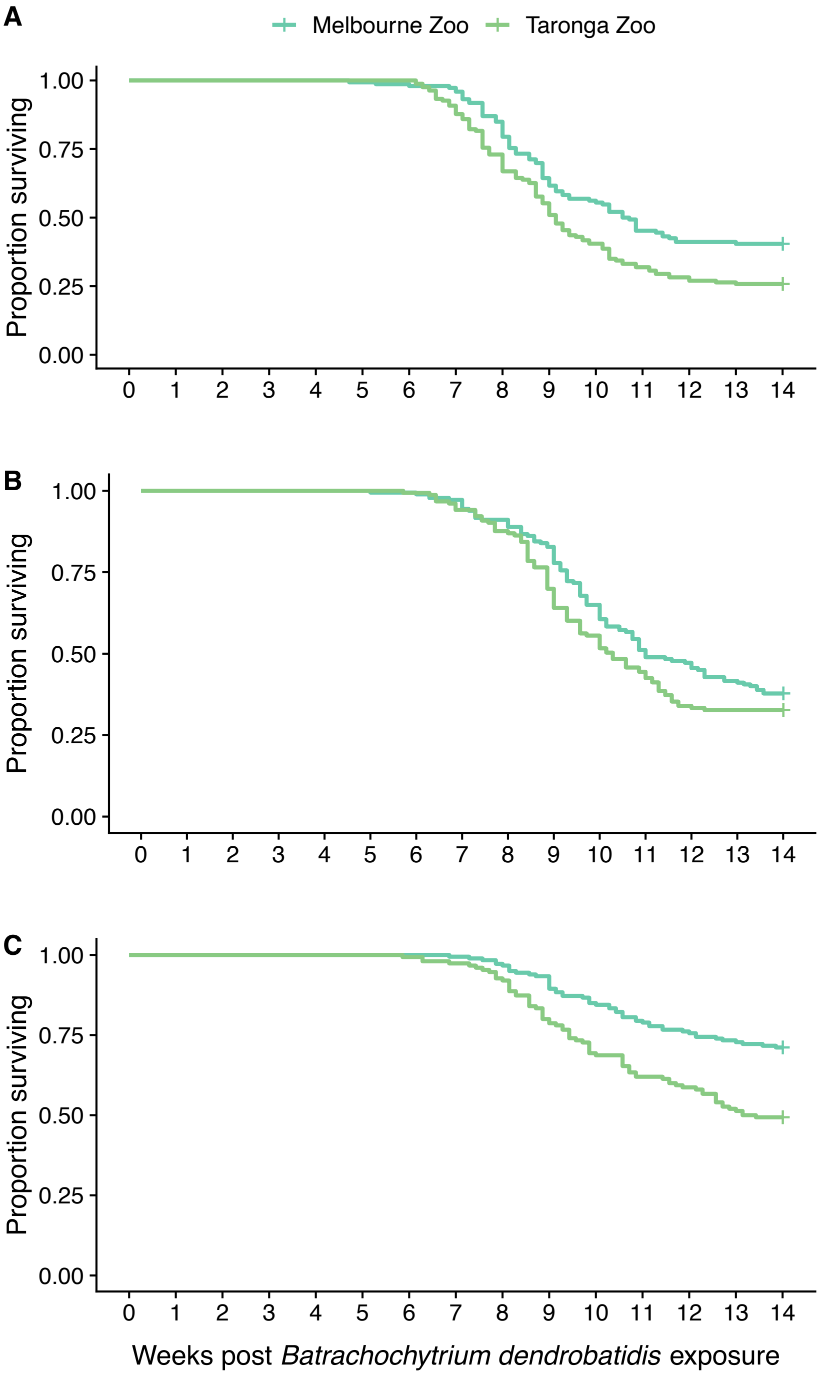


**Figure S2.** Survival outcomes of *P. corroboree* exposed to *Bd* for (**A**) Experiment 1, (**B**) Experiment 2, and (**C**) Experiment 3, by the respective zoo where they were bred and raised.

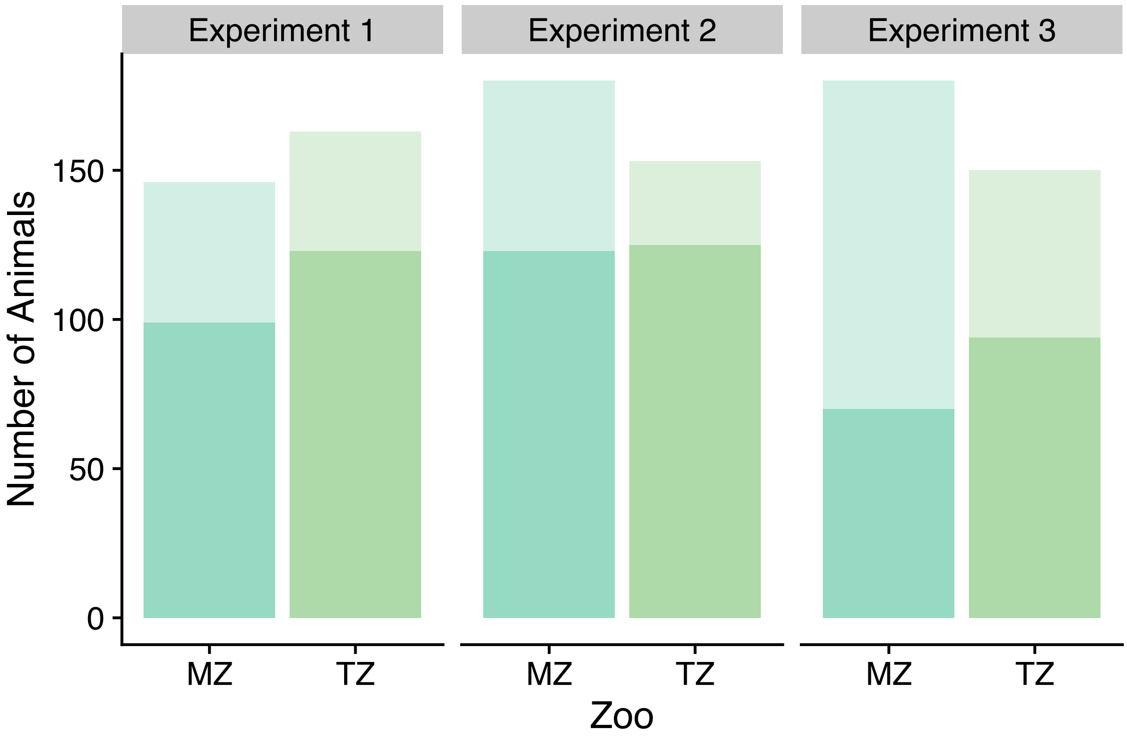


**Figure S3.** Infection status of *P. corroboree* categorised by the zoo where they were bred and raised–Melbourne Zoo (MZ) or Taronga Zoo (TZ)–for each experiment. *Bd*-positive frogs are depicted in full colour, while *Bd*-negative frogs are displayed with reduce opacity.

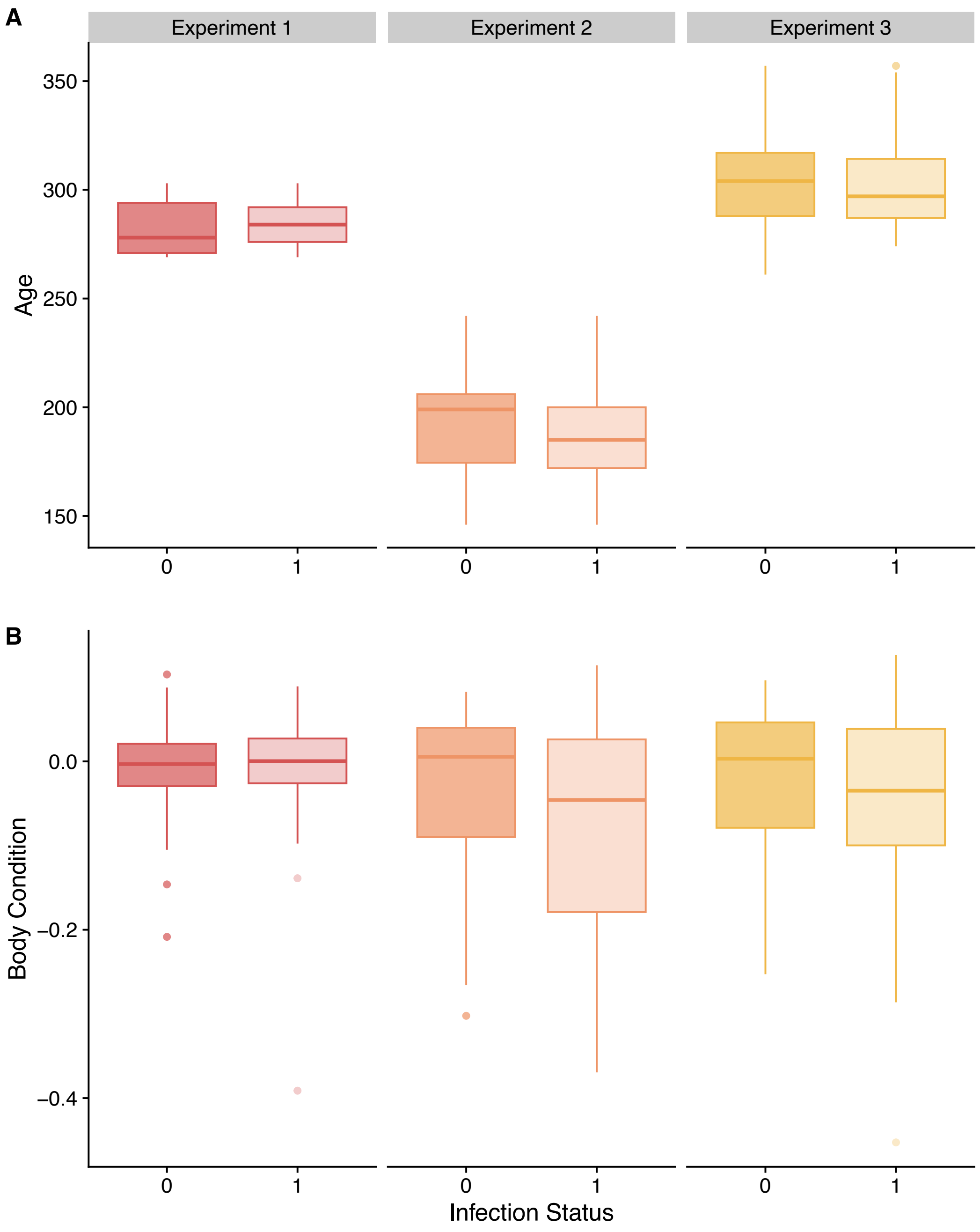


**Figure S4.** Infection status of *P. corroboree* categorised as *Bd*-negative (0) or *Bd*-positive (1), showing (**A**) age in days, and (**B**) body condition across the three experiments.

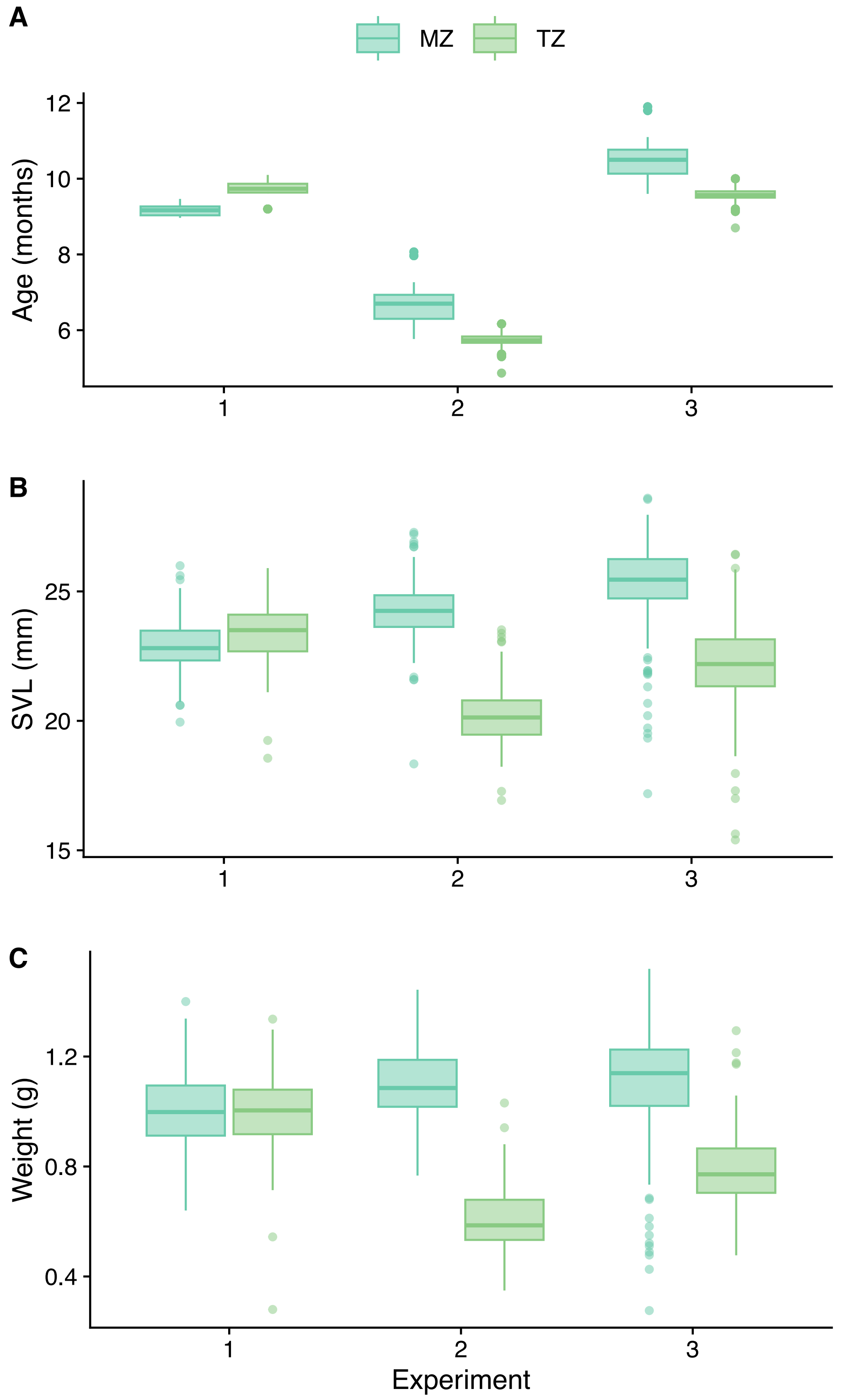


**Figure S5.** Comparison of (**A**) age; (**B**) snout-vent length (SVL); and (**C**) weight of the 972 *P. corroboree* at exposure, grouped by the zoo where they were bred and raised–Melbourne Zoo (MZ) or Taronga Zoo (TZ)–across the three experiments.
